# Epidemiological dynamics of viral infection in a marine picoeukaryote

**DOI:** 10.1101/2021.03.23.436719

**Authors:** Luisa Listmann, Sarah Heath, Pedro F. Vale, C. Elisa Schaum, Sinead Collins

## Abstract

*Ostreococcus tauri* is a ubiquitous marine pico-eukaryote that is susceptible to lysis upon infection by its species specific *Ostreococcus tauri* viruses (OtVs). In natural populations of *O. tauri*, costs of resistance are usually invoked to explain the persistence or reappearance of susceptible individuals in resistant populations. Given the low costs of resistance measured in laboratory experiments with the *O. tauri*/OtV system to date, the question remains of why susceptible individuals persist in the wild at all. Epidemiological models of host and pathogen population dynamics are one useful approach to understand the conditions that can allow the coexistence of susceptible and resistant hosts. We used a SIR (Susceptible-Infected-Resistant) model to investigate epidemiological dynamics under different laboratory culturing regimes that are commonly used in the *O.tauri*/OtV system. When taking into account serial transfer (i.e. batchcycle lengths) and dilution rates as well as different resistance costs, our model predicts that no susceptible cells should be detected under any of the simulated conditions – this is consistent with laboratory findings. We thus considered an alternative model that is not used in laboratory experiments, but which incorporates one key process in natural populations: host populations are periodically re-seeded with new infective viruses. In this model, susceptible individuals re-occurred in the population, despite low costs of resistance. This suggests that periodic attack by new viruses, rather than (or in addition to) costs of resistance, may explain the high proportion of susceptible hosts in natural populations, and underlie the discrepancy between laboratory studies and observations of fresh isolates.

**Importance:** In natural samples of *Ostreococcus* sp. and its associated viruses, susceptible hosts are common. However, in laboratory experiments, fully resistant host populations readily and irreversibly evolve. Laboratory experiments are powerful methods for studying process because they offer a stripped-down simplification of a complex system, but this simplification may be an o*ver*simplification for some questions. For example, laboratory and field systems of marine microbes and their viruses differ in population sizes and dynamics, mixing or migration rates, and species diversity, all of which can dramatically alter process outcomes. We demonstrate the utility of using epidemiological models to explore experimental design and to understand mechanisms underlying host-virus population dynamics. We highlight that such models can be used to form strong, testable hypotheses about which key elements of natural systems need to be included in laboratory systems to make them simplified, rather than oversimplified, versions of the processes we use them to study.

## 1. Introduction

Surveys of biological diversity over the past decades have shown that viruses infecting marine phytoplankton are one of the most numerically important biological entities on the planet (Suttle 2005). Given their high abundance, and in light of several case studies of the possible roles of viruses in structuring marine microbial populations (Schroeder et al. 2003; Suttle 2007), it is well-accepted that marine viruses play a major top-down role in shaping microbial populations in aquatic systems (Suttle 2007; Brussaard 2004; Weitz and Wilhelm 2012), which can then affect aquatic food webs and nutrient cycles. For example, phytoplankton, which include eukaryotic marine primary producer, play an important role in the biological carbon pump when they are grazed or sink as debris to the ocean floor. Here, infections of phytoplankton populations by marine viruses have the potential to alter dynamics of carbon sequestration (Suttle 2007). To understand how host-virus interactions affect host population dynamics and diversity, prokaryotic host-virus laboratory systems have been developed and produced multiple examples of how viruses can drive evolution in their bacterial hosts at the genome, population and community levels (Koskella and Brockhurst 2014; Westra et al. 2019). However, fewer studies exist for eukaryotic microbes such as phytoplankton and their viruses (Short 2012), limiting our ability to integrate host-virus interactions into our understanding of how marine phytoplankton populations are shaped.

We focus on the marine pico-eukaryote *Ostreococcus tauri*. Photosynthetic pico-eukaryotes are less than 3 μm in diameter, and although they are not numerically dominant in oceans, they are important primary producers (Worden, Nolan, and Palenik 2004). *O. tauri* is globally distributed, and is the smallest free-living eukaryote described to date (Courties et al. 1994). Due to its simple cell structure, available genome sequence and ease of culturing, *O. tauri* and its viruses (*Ostreococcus tauri* virus; OtV) have been widely used over the past decade as a model organism for studying marine virus infection (E. Derelle et al. 2006; Clerissi, Desdevises, and Grimsley 2012; Clerissi, Grimsley, Ogata, et al. 2014). To date, *O. tauri* has been found *via* metagenomic analysis in or isolated from several oceanic areas including the Mediterranean, the Aea (Clerissi, Grimsley, Subirana, et al. 2014; Šulčius and Holmfeldt 2016; Zeigler Allen et al. 2017, Listmann et al unpubl data). Where OtVs have been isolated and infections investigated, studies found OtVs to be mainly species specific and readily able to infect hosts (Clerissi, Desdevises, and Grimsley 2012). This indicates that in natural populations susceptible cells (S) are probably present at high frequencies whereas resistant (R) lineages are present at low frequencies, which implies that host resistance is either costly, reversible, or both.

In laboratory studies, resistant (R) cells quickly arise from susceptible (S) populations at a rate of at least one in 1000 (Yau et al. 2016). In addition, viral resistance is both cheap and irreversible once it evolves in laboratory culture, even in the subsequent absence of viruses (Heath et al. 2017; Thomas et al. 2011), which suggests that resistance should be widespread in natural populations. Evidence that both genetic and epigenetic changes are involved in resistance (Yau et al. 2016) could explain why a shift from resistance to susceptibility has not been observed in laboratory populations, in that a genetic constraint could prevent reversion or make it extremely rare. However, if resistance is genetically constrained with little or no cost, this begs the question of why susceptible cells are detected in fresh isolates of *O. tauri* (Clerissi, Desdevises, and Grimsley 2012; Evelyne Derelle et al. 2008) and how S and R *O. tauri* cells coexist in their natural environment.

A possible way to reconcile these data is to consider that the cost of resistance measured in laboratory culture does not reflect the magnitude of that cost in the ocean, but testing this hypothesis experimentally is challenging on a number of levels. A second non-exclusive explanation is that laboratory rearing conditions, which differ drastically from natural conditions, affect host-virus dynamics in ways that could explain this discrepancy.

One useful way to explore how viruses can interact with, and thus affect the biology of their host populations, is through epidemiological models (Heesterbeek et al. 2015; Grassly and Fraser 2008; Lenski 1988; Gandon and Vale 2014). This approach is rarely used in marine (or even phytoplankton) host-virus systems (Middelboe 2000), and here we show that such models can be useful for understanding both the population dynamics of host populations and the relative contributions of resistant (R) and susceptible (S) cells, as well as the population dynamics of the viral strain to which resistance has evolved. The advantage of using models to understand the dynamics of host-virus systems are two-fold: First, they are an important complement to laboratory studies and surveys of natural populations, especially in marine systems, which are relatively difficult to study because of a) the vastness and three-dimensional structure of the marine environment, making it difficult to sample; b) the difficulty of co-culturing marine algae and viruses in the laboratory; and c) the limited population size of experimental populations, where virus infection usually leads to a population crash. Second, in epidemiological models we can explore how aspects of laboratory rearing conditions themselves, such as batch cycle length or dilution rates for culture transfers, affect host-virus interactions. This in turn allows future studies to improve the design and the interpretation of laboratory experiments.

Here, we explore the impacts on the evolution of host resistance of two necessary simplifications of natural environments in laboratory experiments: population dynamics introduced by culturing method, and the lack of immigrants (re-introduction of virus) into host populations. To tackle these questions, we modelled the epidemiological dynamics of this system using a modified SIR (Susceptible-Infected-Resistant) model with different batch cycle lengths or dilution rates and under a range of costs of resistance. We test which magnitude of resistance costs can allow the coexistence of susceptible individuals within a mainly resistant population, and the sensitivity of model outputs to reseeding host populations with fresh viral particles. In particular, we answer the following questions: 1) How do rearing conditions such as growth cycles dominated by exponential vs stationary phase (corresponding to bloom vs non-bloom conditions in natural populations) and dilution rate (corresponding to strong or weak mixing of the water body) and virus re-seeding affect population dynamics of a host-virus system? 2) How does the cost of resistance affect the host-virus system in different rearing conditions? 3) Can we explain the discrepancies in the prevalence of host resistance between natural and laboratory systems?

## 2. The model

### Simulations with one host population and one host-specific virus population

We consider an *O. tauri/OtV* population consisting of susceptible (S), infected (I) and resistant (R) host cells and viruses (V) (Fig. 1). Host cell population size is determined by a birth rate *b* and a death rate *d*; resistant (R) cells experience a cost of resistance which reduces the birth rate by a constant rate *c*. All populations experience regulated growth determined by a carrying capacity *k*. We assume that the starting population already consists of susceptible and resistant cells, with susceptible cells present at a proportion of 100000:1, which is at the minimum of resistant cells as measured previously (Yau et al. 2016; 2018). Infectious OtV viruses are also present in the starting population at a proportion of 100 times less than the host population, as determined empirically in the laboratory (Heath SE., unpublished data). This model assumes the population is well mixed and that host *O. tauri* cells and OtVs are distributed homogenously and come into contact at random. The rate at which an encounter between a host cell and an OtV results in a successful infection is therefore proportional to the densities of the hosts and viruses and the infection rate (*β*. Successful infection of susceptible cells usually results in cell death via lysis (*λ*) with a burst size (δ) of 25 viruses per lysed cell (Evelyne Derelle et al. 2008). However, some infected cells can become resistant at rate *α*. This is in line with previous observations that resistant cells arise spontaneously in *O. tauri* populations exposed to OtV (Thomas et al. 2011). Importantly, we know that resistant populations persist over at least 200 generations in the absence of OtV (Heath et al. 2017). Thus, resistance is inherited, and resistant cells persist in the population and reproduce. We do not include the possibility of resistant cells reverting to become susceptible, since this has never been observed in the laboratory (Yau et al. 2016; 2018; Heath et al. 2017; Thomas et al. 2011), and it is reasonable to assume that the chromosomal rearrangement involved in resistance poses a genetic constraint that makes reversion extremely rare. It has been observed that OtVs are also able to adsorb to resistant cells in the same way as they do to susceptible cells (*β*RV) (Thomas et al. 2011), but do not cause cell lysis in resistant cells. This adsorption rate is assumed to be dynamic enough not to affect the free virus population available to infect cells over the course of a day. Viruses are also removed from the population via decay of the viral particles (*γ*), which we assume to be constant and proportional to the OtV population density. We performed experiments to measure OtV5 decay rate since this parameter was previously unknown (supplementary material S1). For mathematical tractability, adsorption rate *β*, lysis rate *λ*, burst size *δ* and viral decay rate *γ* are assumed to be constant.

**Figure 1.**
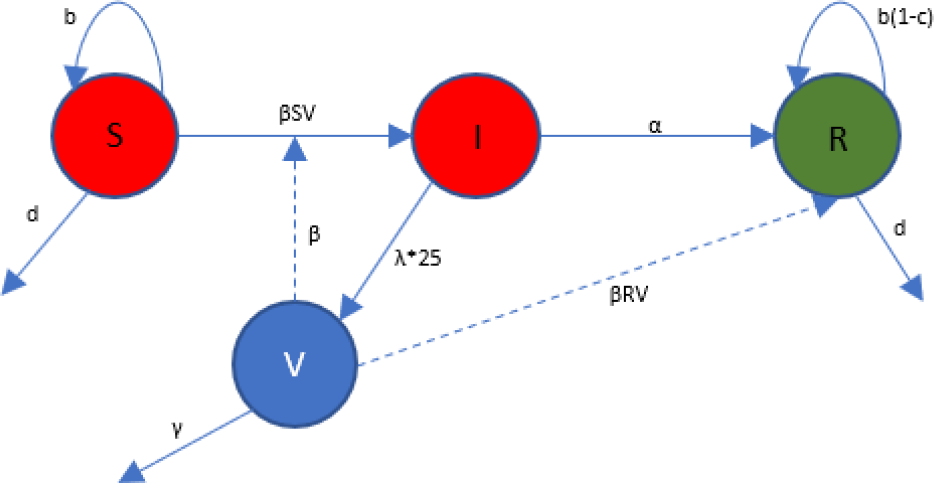
Flow diagram of the model. Circles represent different states of susceptible (S), infected (I) that are also counted into the susceptible population, and resistant (R) states of *O. tauri* cells and viruses (V). Dotted lines indicate effects of viral absorption whereas arrows indicate transitions between states.

The population dynamics for the model can be described by the following ordinary differential equations:

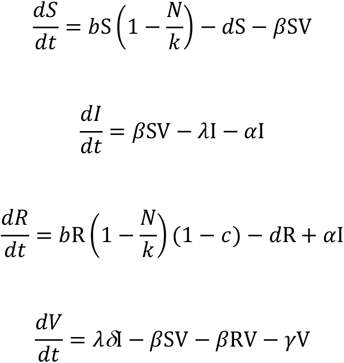

where total cell population size N = S + I + R.

The model simulates a semi-continuous batch culture system where after a number of set integration steps (i.e. batch cycle length=BC) the population was diluted. In the remainder of this study, we will refer to the number of integration steps that elapse between dilutions as ‘transfer rates’. The transfer rates varied between 2 days and 12 days. In addition, we used different quantities of the population for each transfer (i.e. different dilutions) that varied between 0.1 (10% transferred from one batch cycle to the next) and 0.9 (90% transferred, respectively). This allowed to test for the effect of changes to fundamental features of rearing conditions (how often cultures are transferred, and what proportion of the population is transferred) of the host-virus system. The model was then run for a length of 160 model days (integration steps) which led to a different number of maximum batch cycles between 16 (for long batch cycle lengths) and 80 (for short batch cycle lengths). The growth rate we used in the model was 1 which means that the maximum number of generations that a population has gone through was 160 when the model population did not reach stationary phase. Any longer runs would not be comparable to most published laboratory or natural observations in this system. Also, since evolutionary processes readily occur in laboratory experiments on this timescale (Koskella and Brockhurst 2014; Collins 2011), longer timescales were not explored here.

After investigating how rearing conditions affected the dynamics of host resistance assuming that resistance had no cost, we tested how a range of costs of resistance (between 0 and 1, with 0 referring to no cost, and 1 referring to death due to evolution of resistance) affected the modelled populations and the contributions of resistant (R) vs susceptible (S) host cells.

### Extended simulations with the addition of new virus particles at an increasing rate

Due to continuous mixing of the water column under natural conditions and strong host specificity, a resistant host cell can rapidly encounter new virus genotypes that may be infective, despite its resistance to previously encountered virus genotypes. We therefore extended our model such that at specific intervals we added a new virus population that was able to infect resistant host cells, and as such turns them into susceptible cells again (see supplementary material S1 for differential equations). The extended model contained a new susceptible host population (S2), a new infected population (I2) and a new virus population (V2) (Fig. S2 for extended model). We increased the rate of re-seeding the model population with new viruses as follows: The model was set up such that after either 1, 2, 4, or 8 batch cycles, the resistant host cells came into contact with a new virus (V2) to which they were susceptible. This turned them into susceptible cells (S2) that in turn could then become infected (I2). In the extended model we also tested how a range of costs of resistance (between 0 and 1) affected the modelled populations and the contributions of resistant vs. susceptible cells to the whole host population.

We performed all model simulations and approximations of ordinary equations using lsoda (Euler forward solution) in the deSolve package in R (version 3.3.1). Parameter values for the model are defined in Table 1. The procedure of how data were extracted from the simulations and brought together to be shown in the results, is described in detail in the supplementary material S1.

**Table1:** Parameter input to SIR-model

| Parameter | Description | Value | Reference |
| --- | --- | --- | --- |
| $B$ | Birth | 1 division per day | Heath & Collins 2016 |
| $D$ | Death | 0.05 per cell per day | Estimate |
| $\beta$ | Infection rate | 0.35 per cell per day | Derelle *et al.* 2008 |
| $\lambda$ | Lysis rate | 0.99 per cell per day | Estimated based on Derelle *et al.* 2008 |
| $\alpha$ | Conversion rate of I to R | 0.01 per cell per day | Yau *et al.* 2016 |
| $\Gamma$ | Virus decay rate | 0.01 per virus per day | This study |
| $\delta$ | Burst size | 25 per cell day | Derelle *et al.* 2008 |
| $C$ | Cost of resistance | 0-1 | This study |
| $K$ | Carrying capacity | 1 000 000 | This study |
| $BC$ | Batch cycle length | 2-12 days | This study |
| $Dil$ | Dilution rate | 0.1-0.9 | This study |

## 3. Results

### Effect of transfer and dilution rate

In the first round of simulations assuming no cost of resistance, we tested how the transfer and dilution rates affected the host population and determined the conditions where the growth of host populations would remain in exponential phase or reach stationary phase in our simulation. We found that with higher dilution rates (fewer cells transferred), the host populations remained largely in exponential growth, which is in line with expectations from laboratory experiments. This was, however, only the case for fast transfer rates. Specifically, the longer the simulated batch cycle length, the more likely it was that the host populations reached stationary phase (Fig 2.), as expected. Based on these first explorations, we decided to run the subsequent simulations to test the effect of cost of resistance on the composition of host populations in all dilution scenarios and with batch cycle lengths of 4 and 10 days. We used all dilution scenarios to test how the loss of individuals during batch cycle transfers affected the model outcome. The different batch cycle lengths allowed the population to either remain in exponential growth (BC4) for the majority of the simulation, or to reach stationary phase (BC10) early in the simulation, respectively.

**Figure 2.**
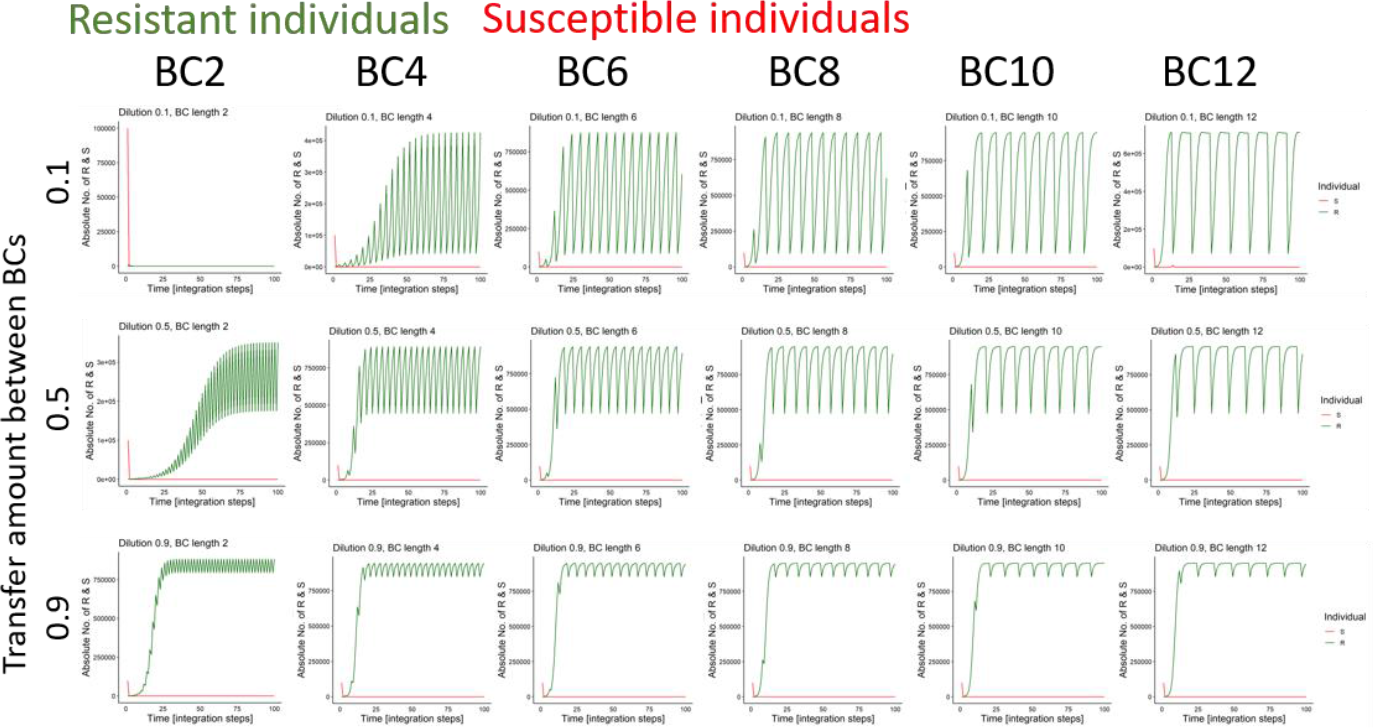
Here the effect of transfer amount (0.1 to 0.9 from one batch cycle to another, with 0.1 indicating a transfer of 10% and 0.9 indicating 90% of the model culture transferred, respectively) (top to bottom) and batch cycle lengths (BC) (left to right) are shown over the model simulations. The number of resistant cells are depicted in green whereas the number of susceptible individuals are depicted in red.

### Effect of cost of resistance on several modelled populations

#### Simulations with one host population and one host-specific virus population

We modelled how different costs of resistance affected the contribution of susceptible (S) and resistant (R) individuals to the host population. In the beginning of simulations (i.e. within one batch cycle) with fast transfer rates (BC 4, Fig. 3a) susceptible individuals were present in host populations. However, susceptible hosts were rapidly lost due to conversion to resistant individuals (Fig 3b-d) and did not change with different costs for resistance. Also, in the rest of the simulations (up to 160 integration steps) we found that including different costs of resistance did not affect the presence of susceptible individuals in the population. Instead, the cost of resistance affected the interaction of the stability of the resistant population and the culturing conditions. Specifically, increasing dilution rates (Figure 3, going from top to bottom in each panel) and increasing the cost of resistance (Figure 3, going from left to right in each panel) led to more experimental populations that collapsed (had a population size below 1000 cells). In addition, the longer the simulations were run (Figure 3, going from left to right panel), the more occasions there were where population collapse happened.

**Figure 3.**
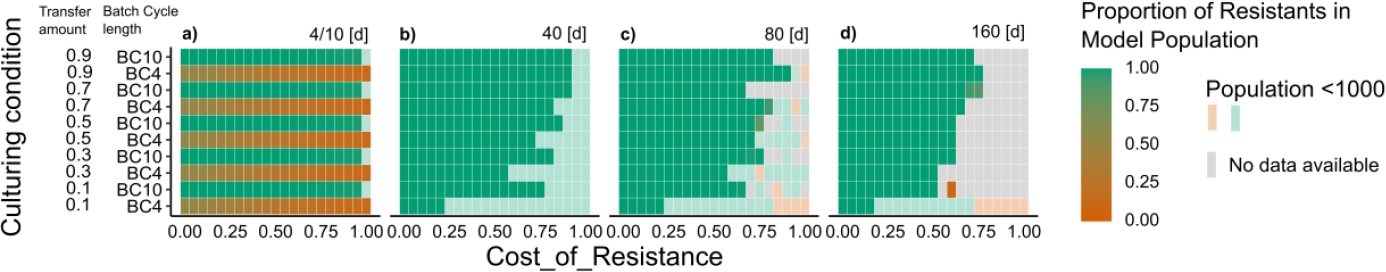
The relative proportion of resistant individuals in the experimental population (green) are shown at the end of the first batch cycle (a) and after 40-160 (b-c) days of modelling run. We note that the number of batch cycles (i.e. transfers) vary for the number of experiments between 4-16 batch cycles when the length is 10 days and between 10 to 80 batch cycles when the length is 4 days. On the x-axis the different modelled costs of resistance are depicted whereas on the y-axis the different culturing conditions are depicted. The colours depict the proportion of resistant (green) vs. susceptible (red) cells in the model populations. When the model population was lower than 1000 individuals the population was defined as “collapsed”.

#### Extended simulations with addition of new virus at an increasing rate

In the extended “virus re-seeding” version of the model, adding new viruses that were able to infect resistant host cells resulted in instances (i.e. simulations) where susceptible cells were present. This in itself is not surprising, but we also found that in the extended model the host populations were less stable. Populations became less stable with i) increasing the rate of re-seeding with viruses (Fig. 4 going from top to bottom), ii) increasing costs of resistance (Fig. 4 going from left to right in each panel), increasing the dilution and transfer rate (Fig. 4 going from top to bottom in each panel). In addition, in approximately 10% of the modelled cases, we found that susceptible cells re-grew in the host populations. That being said, the populations went extinct shortly after susceptible cells regrew. Due to the rapid infection of susceptible cells by the new viruses present in the population, there was not enough time for conversion to resistant cells to stabilize the population (Supplementary Material 2).

**Figure 4.**
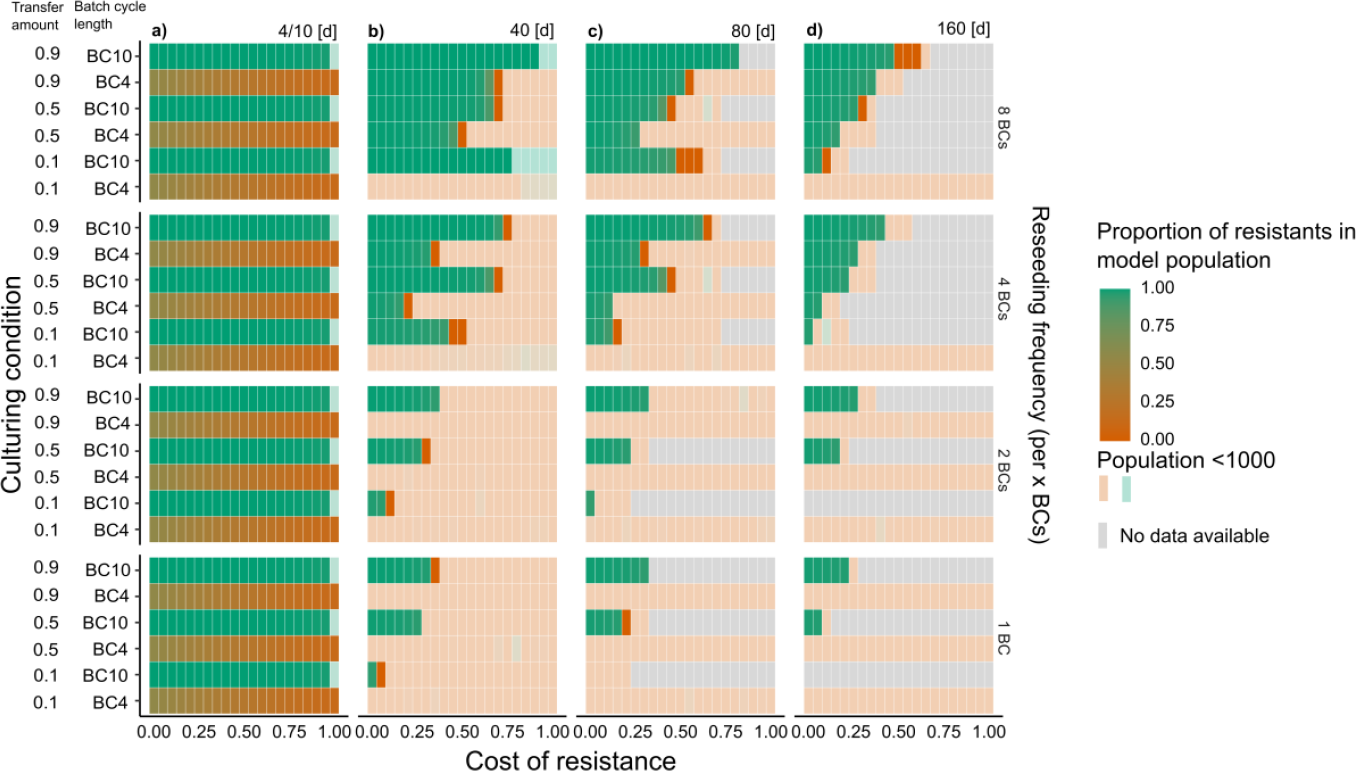
The relative proportion of resistant individuals in the experimental population (green) are shown at the end of the first batch cycle (a) and after 40-160 days (c-d) of modelling run. We note that the number of batch cycles (i.e. transfers) vary for the number of experiments between 4-16 batch cycles when the length is 10 days and between 10 to 80 batch cycles when the length is 4 days. On the x-axis the different modelled costs of resistance are depicted whereas on the y-axis the different culturing conditions are depicted. Going from the top to bottom row the modelling results with increasing virus re-seeding frequency are shown. The colours depict the proportion of resistant (green) vs susceptible (red) cells in the model populations. When the model population was lower than 1000 individuals the population was defined as ‘‘collapsed”.

## 4. Discussion

Mathematical studies of microbial host-virus populations in the oceans extend previously established models, for example by adding a viral component to typical NPZ (nutrient-phytoplankton-zooplankton) food web models (e.g. Weitz et al. 2015) or by manipulating parameters in the Kill the Winner model (Thingstad 2000; Knowles et al. 2016). By using a SIR model, we were able to include the infected proportion of the host population, which is important in showing that *O. tauri* cells have two possible fates following virus infection (lysis or resistance). This allowed us to investigate the fates of host cells under different scenarios: a range of culturing conditions, costs of resistance and different introduction rates of new viruses into the systems.

Our modelling experiments show that culturing conditions can drastically affect experimental outcomes of host-virus dynamics with respect to populations’ resilience to crashes and existence of resistant or susceptible hosts. Specifically, we showed that reaching stationary phase vs staying in exponential phase due to different transfer and dilution rates drastically altered the stability of the modelled populations. SIR models such as this one are thus a useful tool for the design of laboratory host-virus experiments that use batch cultures.

A general explanation for persistence of susceptible hosts within a population is that resistance to viruses is costly, and trades off with growth or reproduction (Antonovics et al. 1994; Sheldon and Verhulst 1996). When viruses are present in the environment, resistance is advantageous and selected for; however, in the absence of viruses, susceptible cells have a fitness advantage if resistance is costly, and should therefore rise in frequency (Lenski, R.E 1988). Previous work has shown costs of resistance to OtV to be very low or undetectable in *O. tauri* in the laboratory under a range of benign or ameliorated conditions (Thomas et al. 2011; Heath et al. 2017). However, since costs of resistance are context-dependent (Meaden, Paszkiewicz, and Koskella 2015; Koskella et al. 2012; Vale et al. 2015; Westra et al. 2015; Labbé, Vale, and Little 2010; Nystrand and Dowling 2020), it is possible that in the ocean, costs exist that have not or cannot be detected in the laboratory. Since we cannot rule out that resistance is in fact costly in natural populations of *O. tauri*, we investigated how varying the costs of resistance affected host populations. In our model, a cost of resistance higher than 0.15 always led to an extinction of the resistant population, which is consistent with laboratory studies failing to detect a cost that is in fact present in natural settings. One explanation for the absence of susceptible cells, when costs of resistance was higher than 0.15, is the rapid infection of any susceptible cells within the starting population. In this case, the only way for the population to avoid extinction at the start of the simulation was indeed to become resistant as quickly as possible. If the cost of being resistant increased, the populations then shortly after went extinct. The transfer and dilution rates in the batch culture system modulated (slowed or speeded) this process, highlighting the context dependence of how costs of resistance affect host population dynamics. Thus, under laboratory conditions, it is not possible to maintain susceptible cells in the system when the evolution of resistance is irreversible. This is due to the uni-directional nature of resistance mutations in this system irrespective or the cost of resistance (Yau et al. 2016).

Another non-exclusive explanation for different results in field observations and modelling is that the dynamics of natural microbe-virus systems are strongly influenced by strain diversity (Agrawal and Lively 2002; Neiman and Fields 2016; Frank 1993), such that some aspects of the natural system cannot be modelled without incorporating diversity. If this is the case, models using single strains of hosts and viruses cannot capture key dynamics that drive the evolution of host resistance, and the maintenance of susceptible hosts, in wild populations. In particular, periodic exposure to new, infective virus strains is important for maintaining susceptible host populations in our simulations. This is in line with current knowledge about the diversity of *Ostreococcus* viruses. For example, Clerissi et al. isolated 40 OtVs that were strain specific, all but two of which came from Mediterranean lagoons (Clerissi, Desdevises, and Grimsley 2012). When re-seeding the modelling experiment with new viruses we then detected instances where susceptible cells were re-detected in the host populations after resistance evolved. However, the frequency of re-detection and relative contribution of susceptible individuals was very low in our model, which is in contrast with the high prevalence of susceptible hosts in natural populations. As before, the frequency of re-detection and relative contribution was context-dependant because of the strong influence of culturing conditions.

Unsurprisingly, we point out that extrapolating from model simulations and laboratory studies to natural environments and interpretations needs to be done with caution. Our current model is not exhaustive and there are more variations that could play a role in how the model populations behave (here we highlight two): 1) different lengths of simulations/experiments and 2) varying numbers of host-virus pairs.

With respect to the length of simulations, we modelled the populations to up to 160 days of experiment. Considering that phytoplankton have fast generation times (in this case up to 1 generation/integration step) and viruses even faster generation times, 160 days of modelled experiment allows reasonable scope for mutations to occur (Elena and Lenski 2003). Fluctuating selection (i.e. the reciprocal evolution of hosts and viruses in response to each other in a negative frequency dependence), causes different rather than increased resistance and infectivity ranges, respectively, with average fitness remaining constant (Buckling and Rainey 2002; Avrani, Schwartz, and Lindell 2012). If fluctuating selection occurs with *O. tauri* and OtVs, this could explain how we do not see decreased fitness in resistant populations. Alternatively, the cost of resistance could be manifested as a limit in the number of virus strains against which a host can be resistant, or as increased susceptibility to other viral strains (Avrani et al. 2011; Marston et al. 2012). However, recent laboratory studies testing resistance specificity in OtV-resistant lines showed an increase in their range of resistance to viruses, probably because resistance to one strain is expected to give rise to resistance to genetically closely related viruses (Yau et al. 2018).

Regarding the point raised with respect to expanding the model: we are only considering a single host – single virus “pair” that is vastly underestimating natural diversity. An individual based modelling approach (Beckmann, Schaum, and Hense 2019; Hinners, Hense, and Kremp 2019) could be better suited to tackling the question of how diversity (either via starting diversity or evolved diversity via mutations) in both host and virus populations more comprehensively affects host-virus dynamics.

## 5. Conclusion

We have described an epidemiological SIR model to investigate *O. tauri*/OtV5 infection dynamics, with a focus on the effect of culturing conditions on the host-virus dynamics. We demonstrate the application of traditional epidemiological models as a useful approach to study marine algae/virus systems and build on data collected from the laboratory and the field.

## Acknowledgements

S.E. Heath was supported by a BBSRC EASTBIO Doctoral Training Programme grant, P. Vale is supported by a Chancellor’s fellowship (University of Edinburgh) and a Branco Weiss fellowship from Society in Science (https://brancoweissfellowship.org/). S. Collins was supported by a Royal Society University Research Fellowship during part of this project…Luisa Listmann and Elisa Schaum are supported by a start-up grant to ES by Universit<u>ät Hamburg</u>.

